## Supplemental Figures S1-S7 for "Folding kinetics of an entangled protein"

### Supplementary Figures

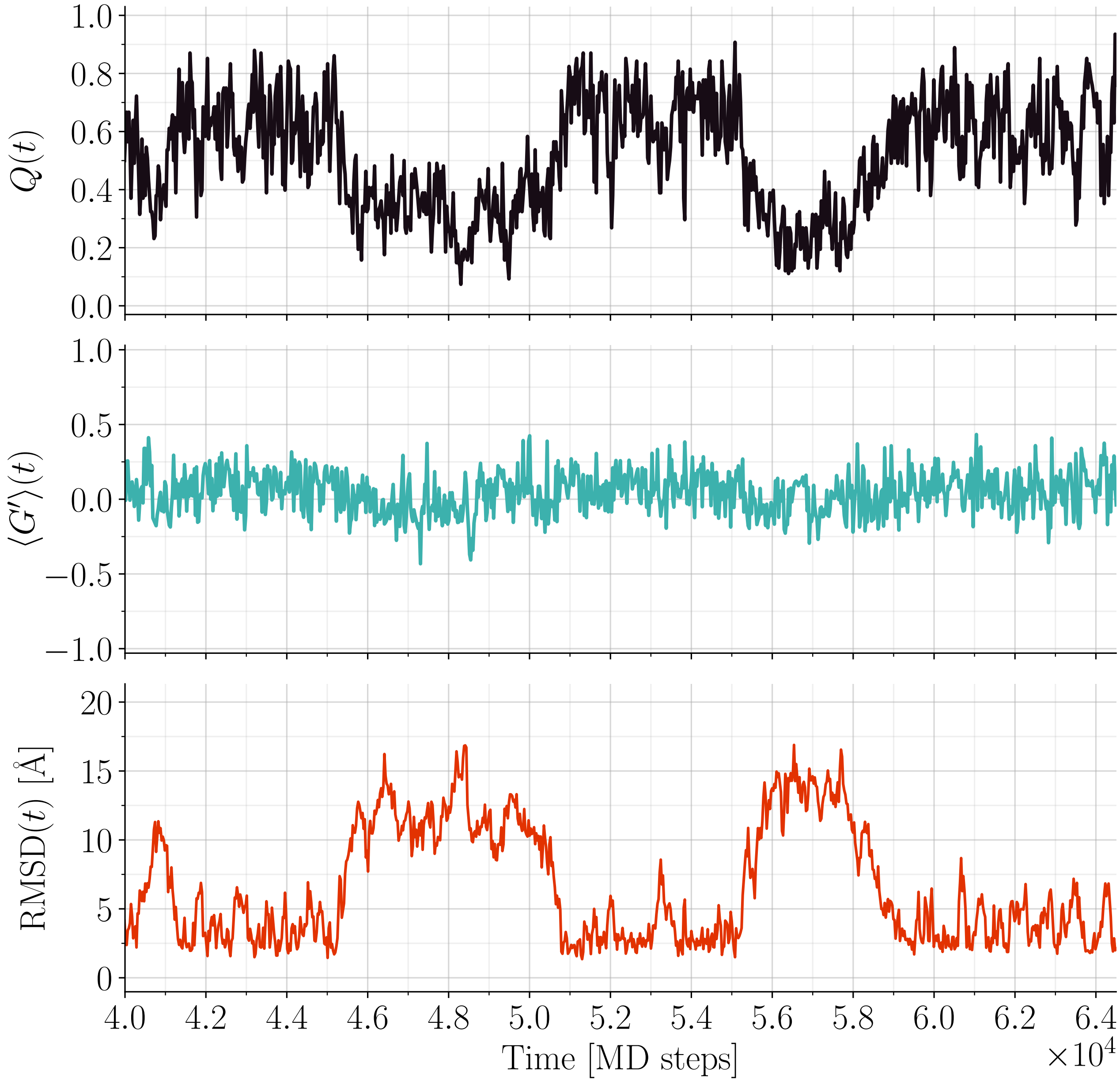


FIG. S1. **Time series from equilibrium simulations for the non-entangled SH3 domain at $T = T_f$.** Upper panel: fraction of native contacts $Q$. Middle panel: entanglement indicator $\langle G' \rangle$. Lower panel: RMSD with the native structure.

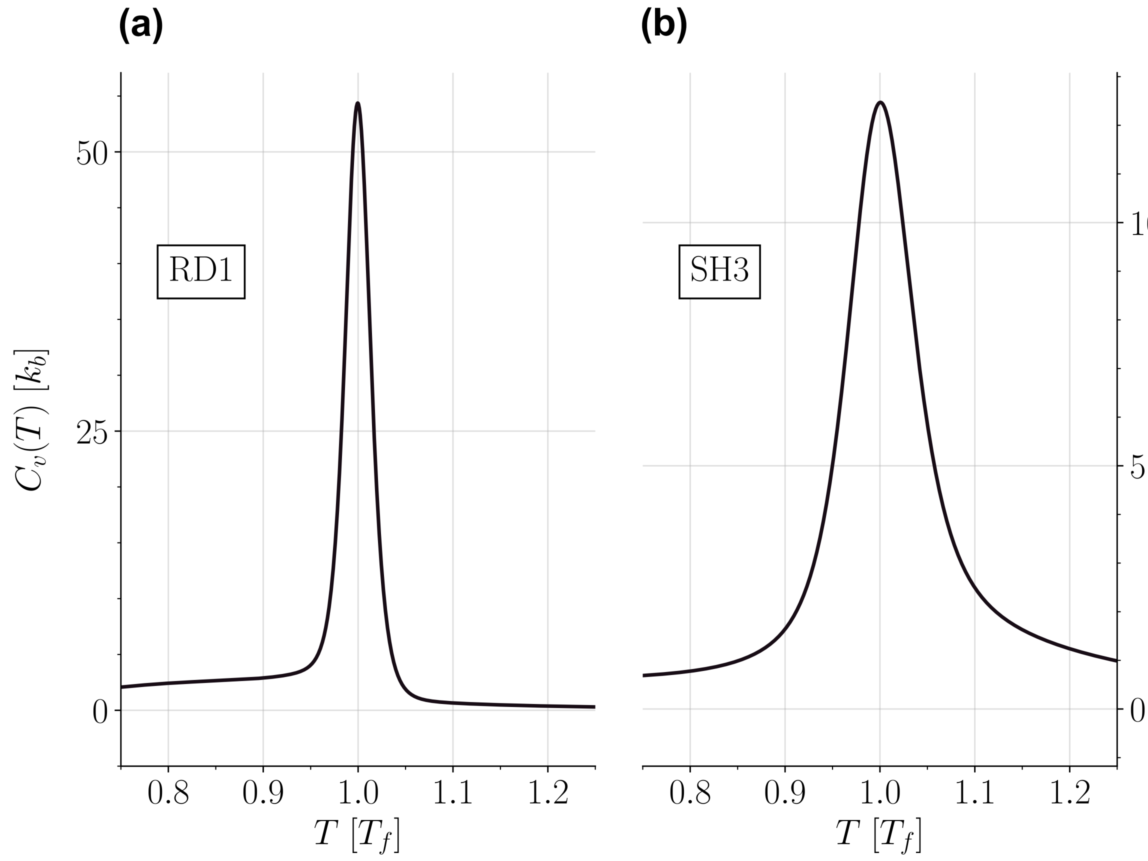


FIG. S2. **Specific heat profiles for the entangled RD1 protein and the non-entangled SH3 domain.** Left panel: RD1 protein. Right panel: SH3 domain. Temperatures are rescaled in both cases by the folding temperature $T_f$ identified at the specific heat peak, which are $T_f^{\mathrm{RD1}} = 1.149 \pm 0.005$ and $T_f^{\mathrm{SH3}} = 1.012 \pm 0.001$.

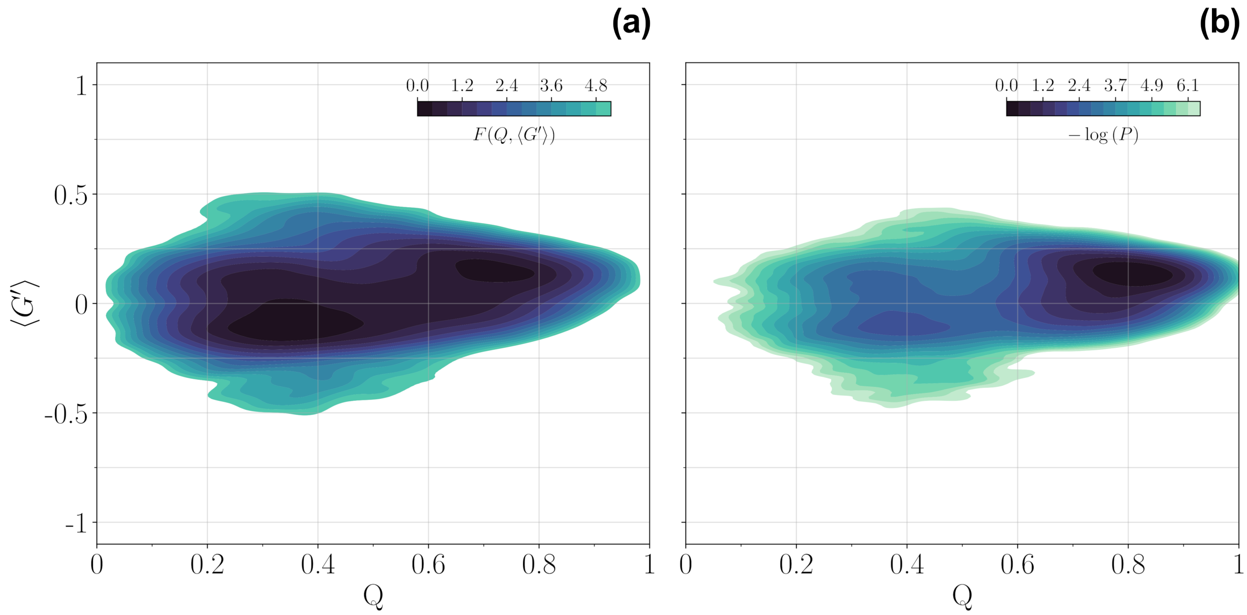


FIG. S3. **Log-scale histogram contour plots in the** $(Q, \langle G' \rangle)$ **plane for the non-entangled SH3 domain.** Histogram negative log-counts are smoothed using KDE (see the Definition of ensembles subsection) and shifted in order for their minimum to be 0. Contour levels and the colour scale are the same in both plots. (a): Data collected from 8 long equilibrium trajectories at $T = T_f$; the contour plot represents a dimensionless free energy surface (b): Data collected from 100 trajectories refolding from an initial unfolded configuration at $T = 0.9T_f$; the unfolded state is populated transiently and the contour plot does *not* represent a free energy surface.

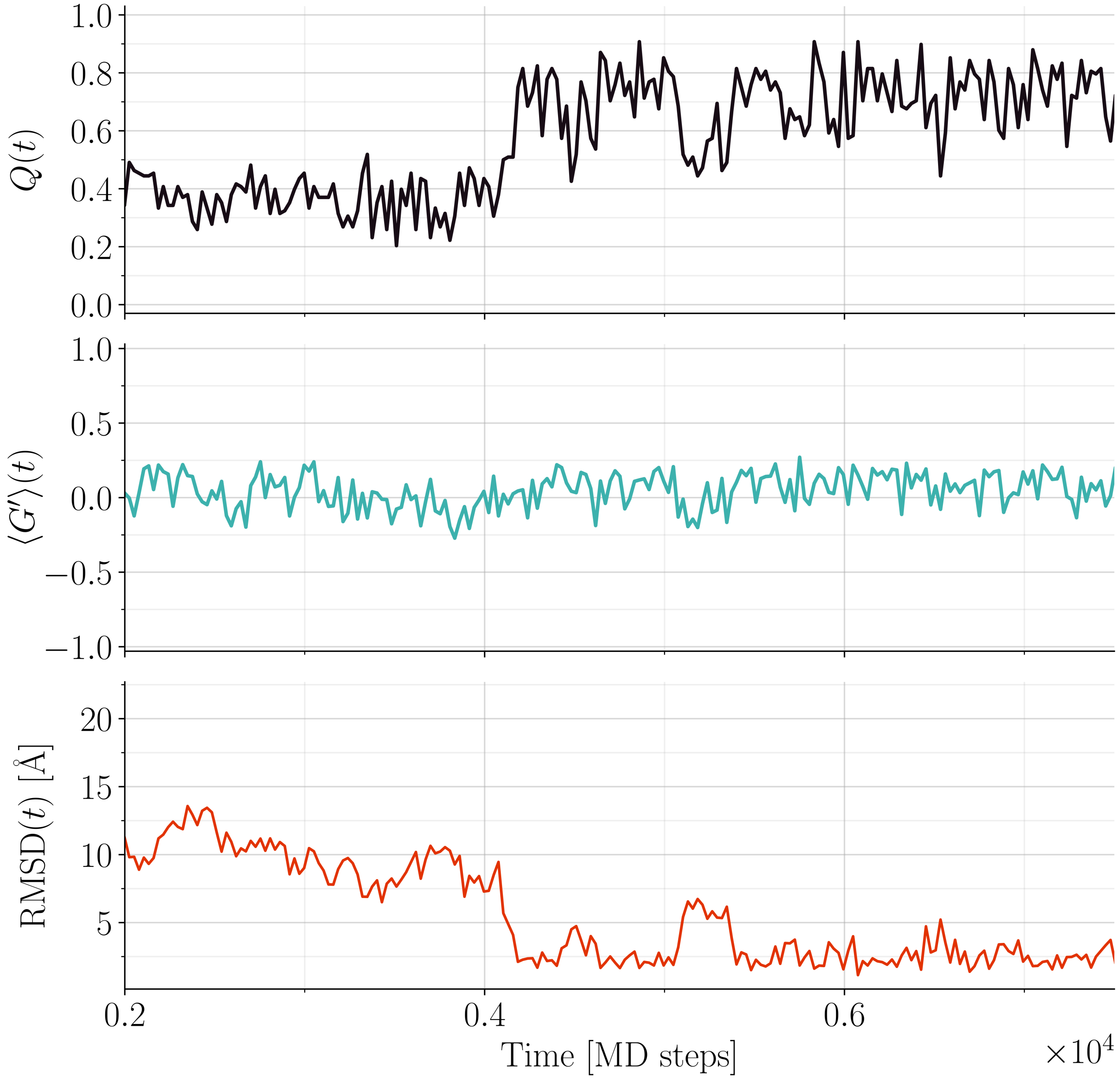


FIG. S4. **Time series from refolding simulations for the non-entangled SH3 domain at $T = 0.9T_f$.** Upper panel: fraction of native contacts $Q$. Middle panel: entanglement indicator $\langle G' \rangle$. Lower panel: RMSD with the native structure.

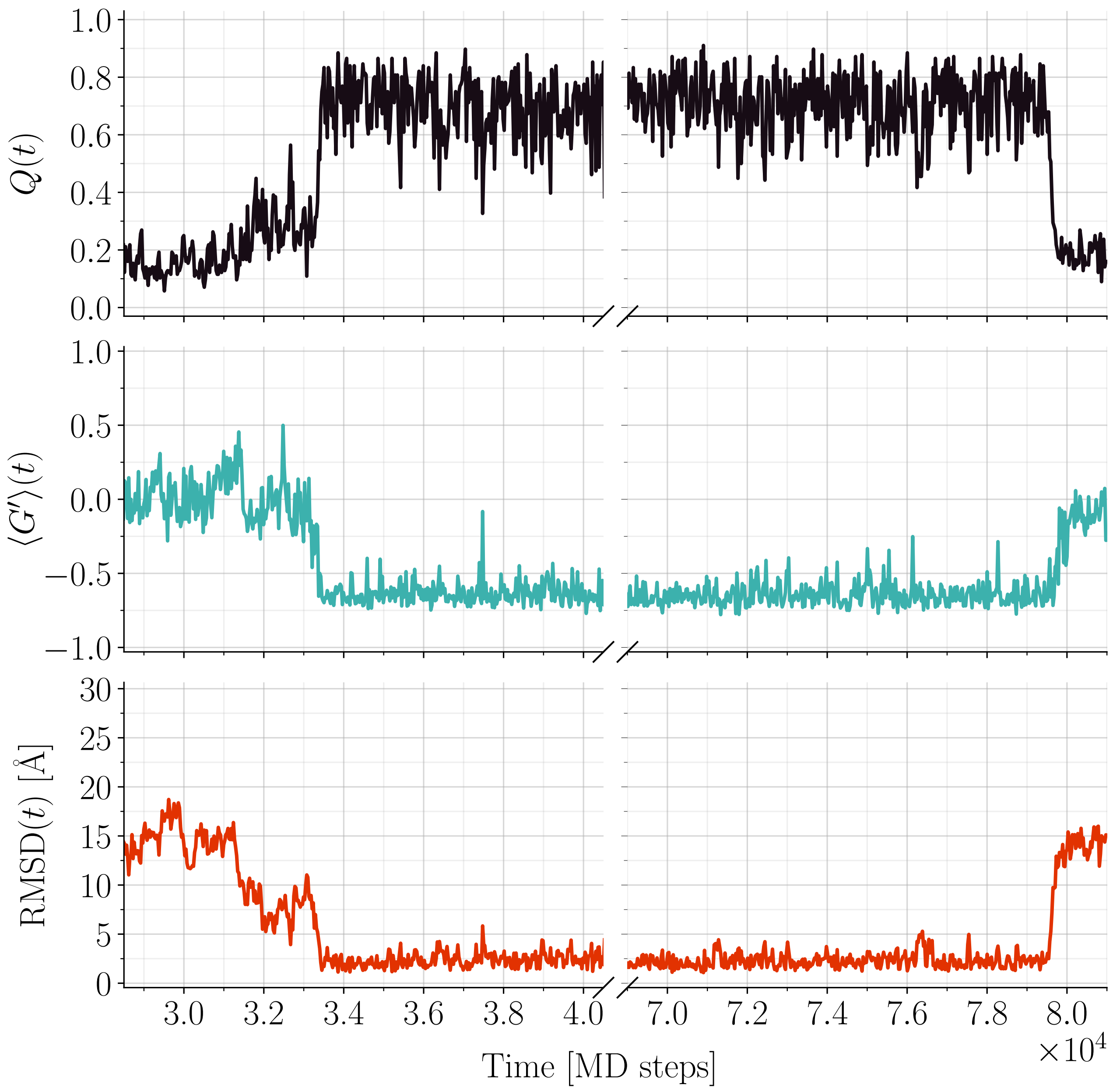


FIG. S5. **Time series from equilibrium simulations for the entangled RD1 protein at $T = T_f$.** Upper panel: fraction of native contacts $Q$. Middle panel: entanglement indicator $\langle G' \rangle$. Lower panel: RMSD with the native structure.

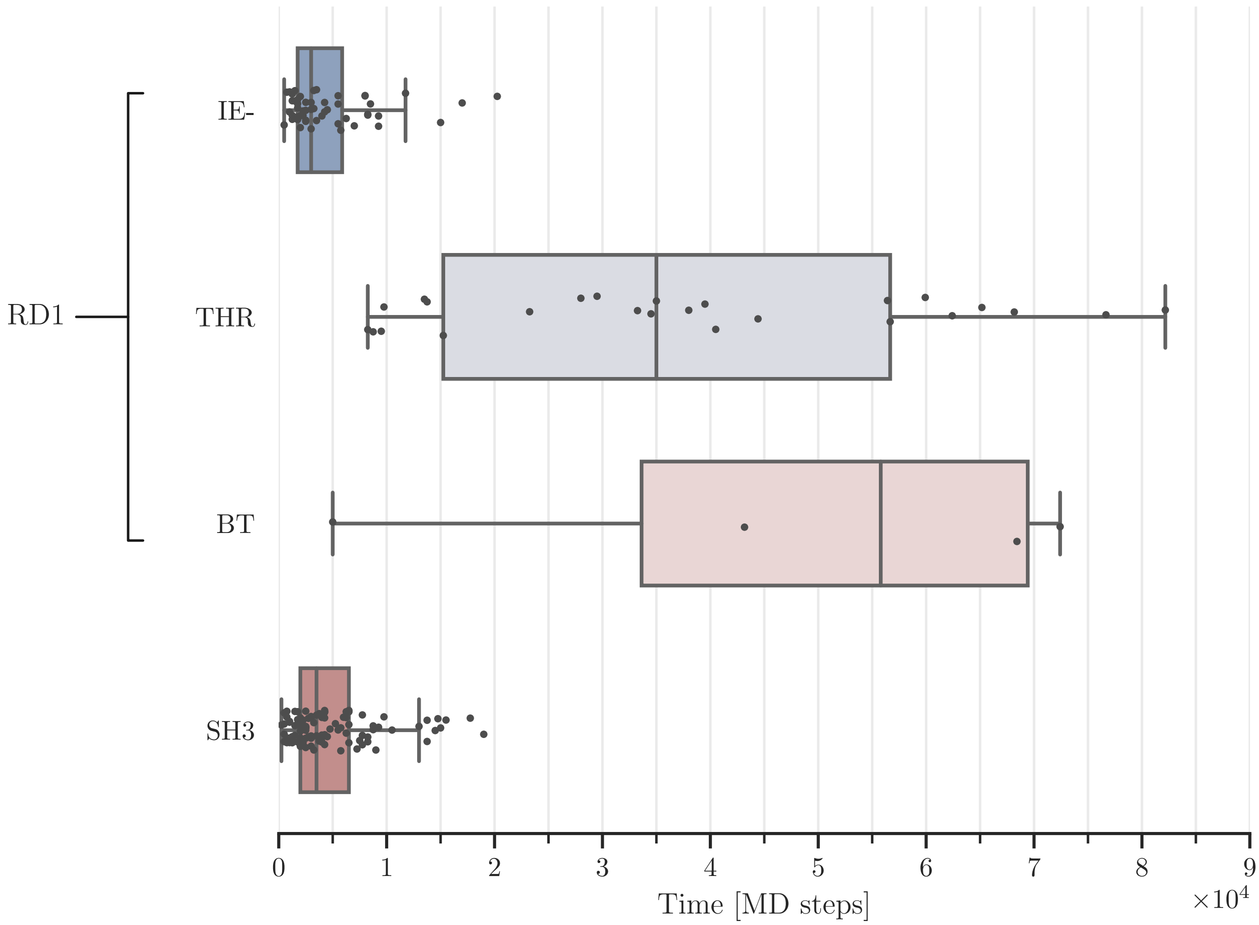


FIG. S6. **Distributions of folding times for the different types of refolding trajectories for the entangled RD1 protein and for the non-entangled SH3 domain.** The 81 trajectories that achieved successful refolding for the RD1 protein are partitioned into the "fast" channel $IE_-$ (52 trajectories going through the short-lived entangled intermediate $IE_-$ that are never trapped in IT), the "threading" channel THR (25 trajectories that fold through a final IT→F transition) and the backtracking channel BT (4 trajectories that finally backtrack to the fast channel after trapping in IT). All the 100 refolding trajectories for the SH3 domain are grouped together. Box-plots show the median, the quartile and the first and ninth decile values.

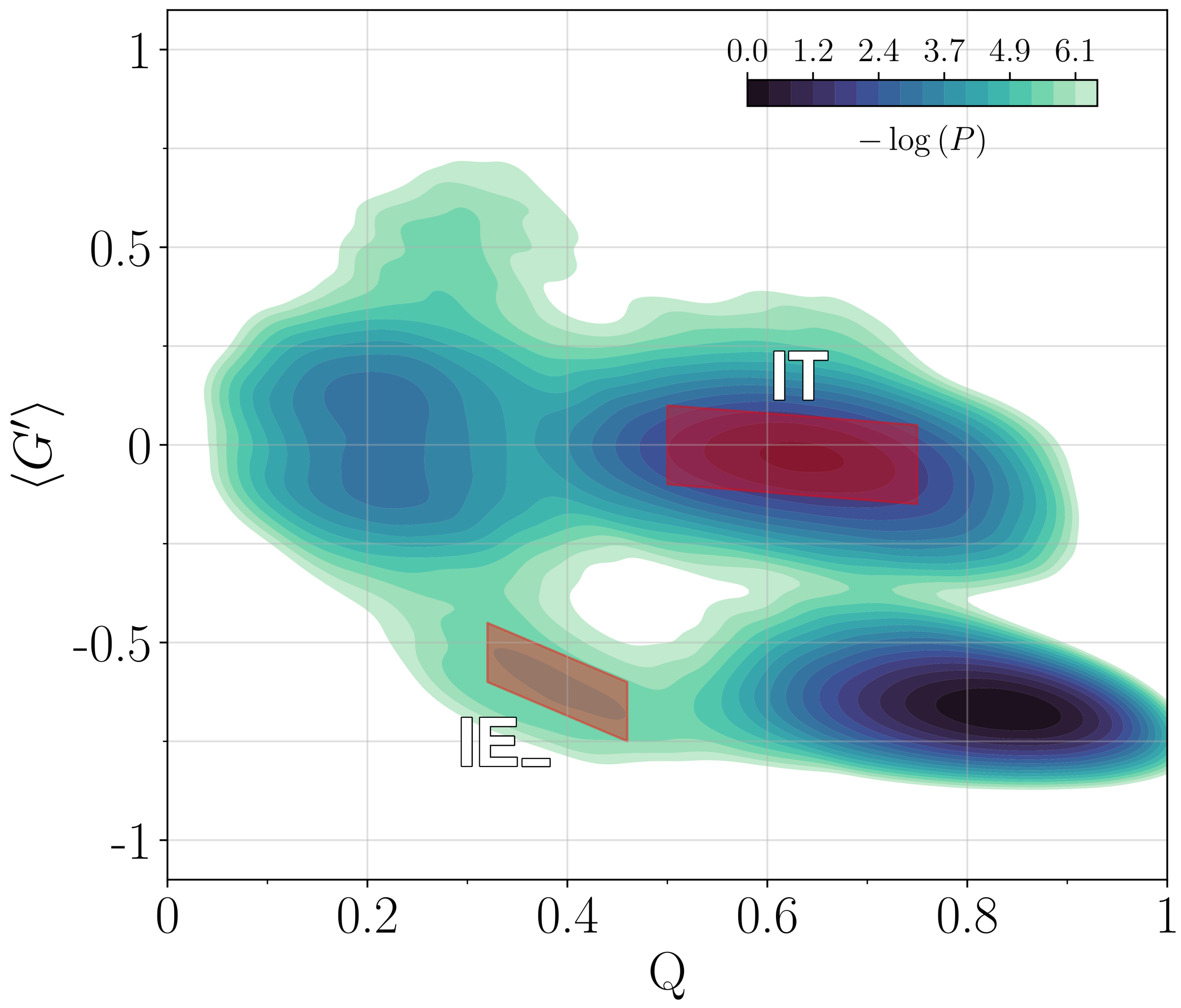


FIG. S7. **Identification of the intermediate state ensembles for the entangled RD1 protein.** Log-scale histogram contour plot in the $(Q, \langle G' \rangle)$ plane from 100 refolding trajectories for the RD1 protein at $T = 0.9T_f$. Colored areas define configurations classified as part of one of the intermediate ensembles. The darker shaded area corresponds to the IT ensemble, while the lighter one to $\mathrm{IE}_-$ ensemble.
